# Hypoxia blunts angiogenic signaling and upregulates the antioxidant system in elephant seal endothelial cells

**DOI:** 10.1101/2023.07.01.547248

**Authors:** Kaitlin N Allen, Julia María Torres-Velarde, Juan Manuel Vazquez, Diana D Moreno-Santillan, Peter H Sudmant, José Pablo Vázquez-Medina

**Affiliations:** Department of Integrative Biology, University of California Berkeley, Berkeley, CA, USA 94720; Center for Computational Biology, University of California Berkeley, Berkeley, CA, USA 94720

**Keywords:** diving, redox, marine mammal, inflammation, glutathione

## Abstract

Elephant seals experience extreme hypoxemia during diving bouts. Similar depletions in oxygen availability characterize pathologies including myocardial infarction and ischemic stroke in humans, but seals manage these repeated episodes without injury. However, the real-time assessment of the molecular changes underlying protection against hypoxic injury in seals remains restricted by their at-sea inaccessibility. Hence, we developed a proliferative arterial endothelial cell culture system to assess the molecular response to prolonged hypoxia. Seal and human cells exposed to 1% O_2_ for up to 6 h demonstrated differential responses to both acute and prolonged hypoxia. Seal cells decouple stabilization of the hypoxia-sensitive transcriptional regulator HIF-1α from angiogenic signaling at both the transcriptional and cellular level. Rapid upregulation of genes involved in the glutathione (GSH) metabolism pathway supported maintenance of GSH pools and increases in intracellular succinate in seal but not human cells during hypoxia exposure. High maximal and spare respiratory capacity in seal cells after hypoxia exposure occurred in concert with increasing mitochondrial branch length and independent from major changes in extracellular acidification rate, suggesting seal cells recover oxidative metabolism without significant glycolytic dependency after hypoxia exposure. In sum, our studies show that in contrast to human cells, seal cells adapt to hypoxia exposure by dampening angiogenic signaling, increasing antioxidant protection, and maintaining mitochondrial morphological integrity and function.

## Introduction

Extreme, repeated bouts of hypoxemia characterize diving in northern elephant seals (*Mirounga angustirostris*) (Elsner et al., 1970; Kerem and Elsner, 1973; Meir et al., 2009; Meir et al., 2013). In the vasculature, hypoxia shifts the endothelium toward a pro-inflammatory state (Janaszak-Jasiecka et al., 2021; Luo et al., 2022; Ten and Pinsky, 2002), priming vessels for increased leukocyte adhesion upon reoxygenation (Granger et al., 1993). In addition, hypoxic changes in mitochondrial metabolism lead to succinate accumulation and subsequent mitochondrial superoxide generation *via* reverse electron transport (Chouchani et al., 2014; Chouchani et al., 2016; Koziel and Jarmuszkiewicz, 2017), though degree of accumulation may modulate these effects (Bundgaard et al., 2023). Reoxygenation reintroduces oxygen to the “primed” system, driving additional oxidant generation from various sources including NADPH oxidases and the electron transport chain. The resulting oxidative damage promotes endothelial dysfunction which compromises both the barrier and vasoregulatory roles of the endothelium (Incalza et al., 2018).

Hypoxia-induced endothelial dysfunction would catastrophically impact the dive response in diving mammals, for whom persistent peripheral vasoconstriction and tight regulation of vascular tone are critical during prolonged submersion (Blix, 2018; Hindle et al., 2019; Zapol et al., 1979). Recent work shows that nitric oxide responsiveness differs between organs and their supplying arteries in Weddell seals (Hindle et al., 2019) and that inflammatory signaling is blunted in Weddell and northern elephant seal blood (Bagchi et al., 2018), suggesting that diving mammals tightly regulate the inflammatory response to mitigate vascular damage during diving. Additionally, marine mammals exhibit a robust antioxidant defense system which likely counteracts the damaging effects of diving-induced oxidant generation. In particular, marine mammals possess high tissue and circulating levels of glutathione (GSH), the most abundant cellular thiol and a crucial cofactor for several antioxidant enzymes (Cantú-Medellín et al., 2011; Forman et al., 2009; Vázquez-Medina et al., 2006; Vázquez-Medina et al., 2007; Vázquez-Medina et al., 2011a; Vázquez-Medina et al., 2011b).

Besides comparative biochemical studies showing that marine mammals exhibit high tissue and circulating levels of GSH, recent work shows duplication events and positive selection on candidate genes involved in GSH metabolism in diving mammals (Foote et al., 2015; Tian et al., 2019; Tian et al., 2021), but the dynamic regulation of the antioxidant system during hypoxia and diving remains unknown. Comprehensive assessments of antioxidant function and immunomodulation remain infeasible during diving due to technological limitations on *in vivo* studies and the inaccessibility of the animals when diving at sea. To this end, we developed a primary cell culture system using arterial endothelial cells derived from expelled placental arteries obtained during elephant seal breeding haulouts. Arteries are generally inaccessible in living marine mammals, and sampling of arterial tissue is constrained to expelled tissues or those derived from necropsy. We then conducted comparative transcriptomic and metabolic analyses to study the molecular mechanisms that drive hypoxemic tolerance in seal endothelial cells. We found that rapid hypoxic stabilization of HIF-1α in elephant seal cells co-occurred with downregulated inflammatory signaling and a highly dynamic transcriptional response promoting vascular homeostasis alongside a blunted migratory response to hypoxia. Moreover, hypoxia increased GSH and succinate levels in seal but not human cells in coordination with enhanced respiratory capacity following hypoxia/reoxygenation. Together, these results highlight the importance of the GSH antioxidant system and mitochondrial function in protecting the vascular endothelium in a diving mammal exposed to extreme oxygen fluctuations.

## Results

### Characterization of elephant seal arterial endothelial cells in primary culture

Primary endothelial cells derived from seal and human placental arteries stained positive for the endothelial marker PECAM-1 (CD31), took up acetylated low-density lipoprotein (Ac-LDL), and spontaneously formed tubes when cultured on a 3D matrix (Figure 1A). CD144 (VE-cadherin) and CD31 mRNA expression was 13-fold greater in seal endothelial cells compared to placental trophoblast cells (CD31: t=5.665, p=0.005; CD144: t=6.501, p=0.003) (Figure 1B). These results confirm the endothelial phenotype of our preparations.

**Figure 1.**
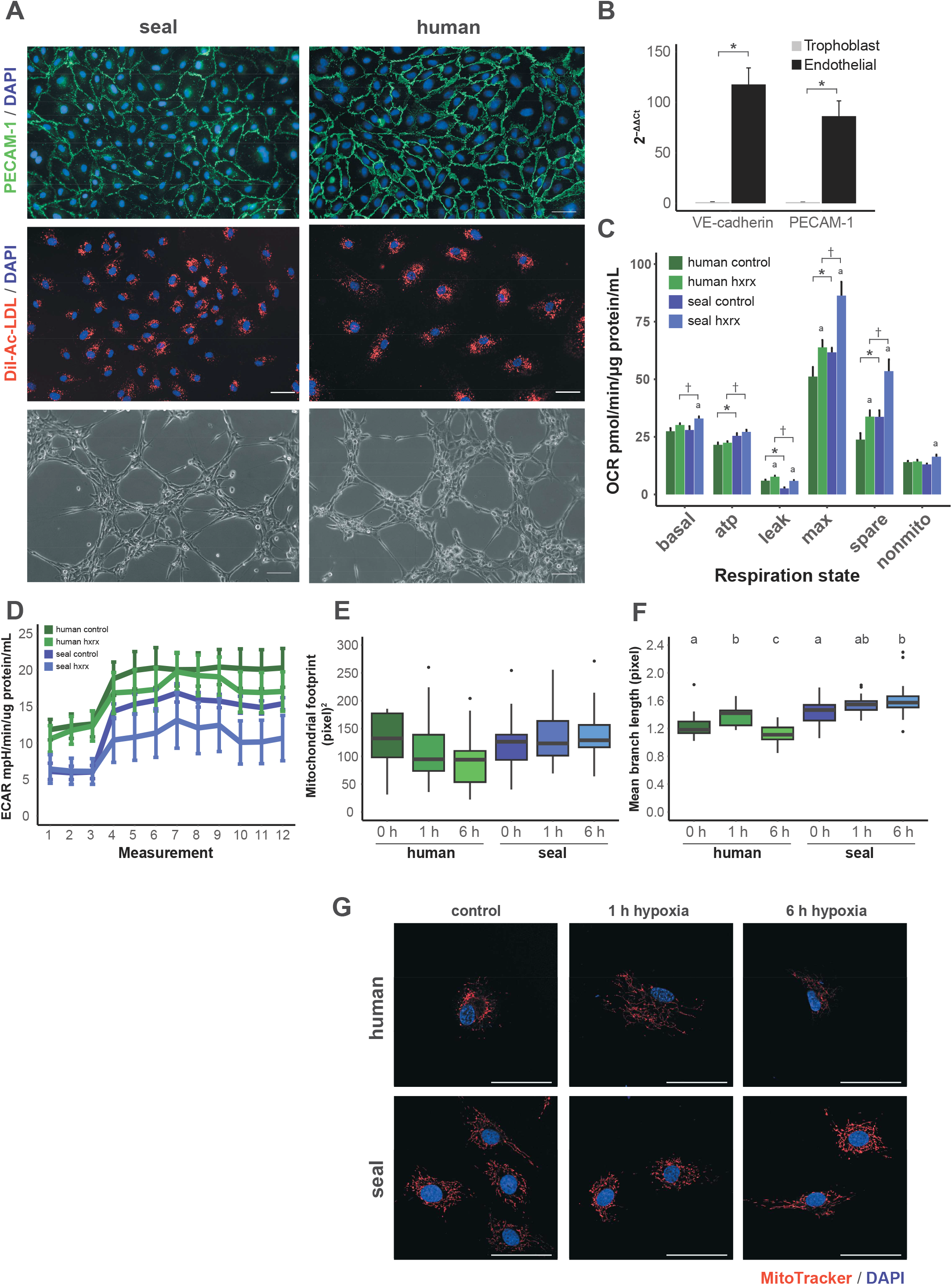
Primary endothelial cells express canonical endothelial markers and modulate cellular respiration in response to hypoxia/reoxygenation. (A) Top: PECAM-1 (CD31; green) staining in seal and human endothelial cells. Middle: DiI-acetylated LDL (red) update by seal and human cells. Nuclei (blue) in top and middle panels are stained with DAPI. Lower: Seal and human endothelial cells spontaneously form tunes in a 3D gel matrix. Scale bar in all images is 50 µm. (B) Relatively expression of the endothelial markers VE-cadherin (CD144) and PECAM-1 (CD31) in seal endothelial compared to placental trophoblast cells, n=3-5. * p<0.05. (C) Mitochondrial function at baseline and after hypoxia/reoxygenation in seal and human cells, n=9. Values are normalized to protein content per well. hxrx, hypoxia/reoxygenation. * p<0.05 between species at baseline. † p<0.05 between species after hypoxia/reoxygenation. Lettering indicates intraspecific changes between baseline and hypoxia/reoxygenation. (D) Extracellular acidification rate (ECAR) in seal and human endothelial cells at baseline and after hypoxia/reoxygenation. (E, F) Mitochondrial footprint (E) and mean mitochondrial network branch length (F) in seal and human endothelial cells after hypoxia/reoxygenation. Hypoxia exposure occurred for 1 or 6 h, followed by 30 min reoxygenation. Lettering indicates intraspecific changes between baseline and hypoxia/reoxygenation. (G) Representative images of human (top) and seal (bottom) endothelial cells stained with MitoTracker Red CMXRos after 0, 1, and 6 h hypoxia followed by 30 min reoxygenation. Scale bar is 50 µm.

Basal oxygen consumption rates did not differ between seal and human cells, though hypoxia/reoxygenation increased basal oxygen consumption only in seal cells. ATP-linked respiration, maximal respiration, and spare respiratory capacity were greater in seal than in human cells at baseline and after hypoxia/reoxygenation despite hypoxia/reoxygenation-induced increases in maximal respiration and spare capacity in both species (Figure 1C). In contrast, human cells displayed greater proton leak than seal cells both at baseline and after hypoxia/reoxygenation, though leak increased in both species following hypoxia/reoxygenation (Figure 1C). While seal cells after hypoxia/reoxygenation displayed the highest overall oxygen consumption rates (Figure 1C), these cells exhibited the lowest extracellular acidification rates, suggesting that glycolytic acidification of the media is not a primary component of these observed increased respiratory capacity (Figure 1D).

We assessed mitochondrial network features to determine whether the observed changes in oxygen consumption may be driven by morphological differences in mitochondria following hypoxia/reoxygenation. Mitochondrial footprint, the total volume of mitochondria in the cell, appeared to decline in human cells after 1 h and 6 h hypoxia followed by 30 min reoxygenation, though this trend was not statistically significant. Mitochondrial branch length increased transiently in human cells after 1 h hypoxia but dropped below baseline levels by 6 h (χ^2^(3)=17.47, p=0.0002). In seal cells, however, hypoxia increased branch length from control to 6 h (χ^2^(3)=9.000, p=0.01). These results suggest that seal cells increase mitochondrial connectivity while maintaining overall volume during hypoxia despite long-term decreases in branching and volume in human cells.

### Hypoxia differentially regulates HIF-1 signaling in seal and human cells

We evaluated the HIF-1-mediated response to sustained hypoxia in seal and human cells. Both human and seal cells remained viable (>99%) throughout a 6 h exposure to 1% O_2_ (data not shown). Hypoxia increased HIF-1α protein levels to a maximum of ∼70-fold over (normoxic) baseline abundance in both species (p=0.005 both species) (Figure 2A,B), though we observed species-specific dynamics. HIF-1α abundance was biphasic in seal cells: an initial major peak within 15 min of hypoxia was followed by a ∼45% decline from 30-120 min, with levels again increasing after 2 h. This late-stage increase co-occurred with an increase in HIF-1α mRNA levels (Figure 2A,C). In contrast, HIF-1α levels did not peak in human cells until 1 h in hypoxia, and abundance declined continuously following this peak. HIF-1α mRNA levels in human cells remained relatively stable through 2 h, followed by a precipitous decline at later timepoints (Figure 2A,C). Together, these data suggest that hypoxia induces rapid and robust stabilization of HIF-1α in seal, but not human, cells.

**Figure 2.**
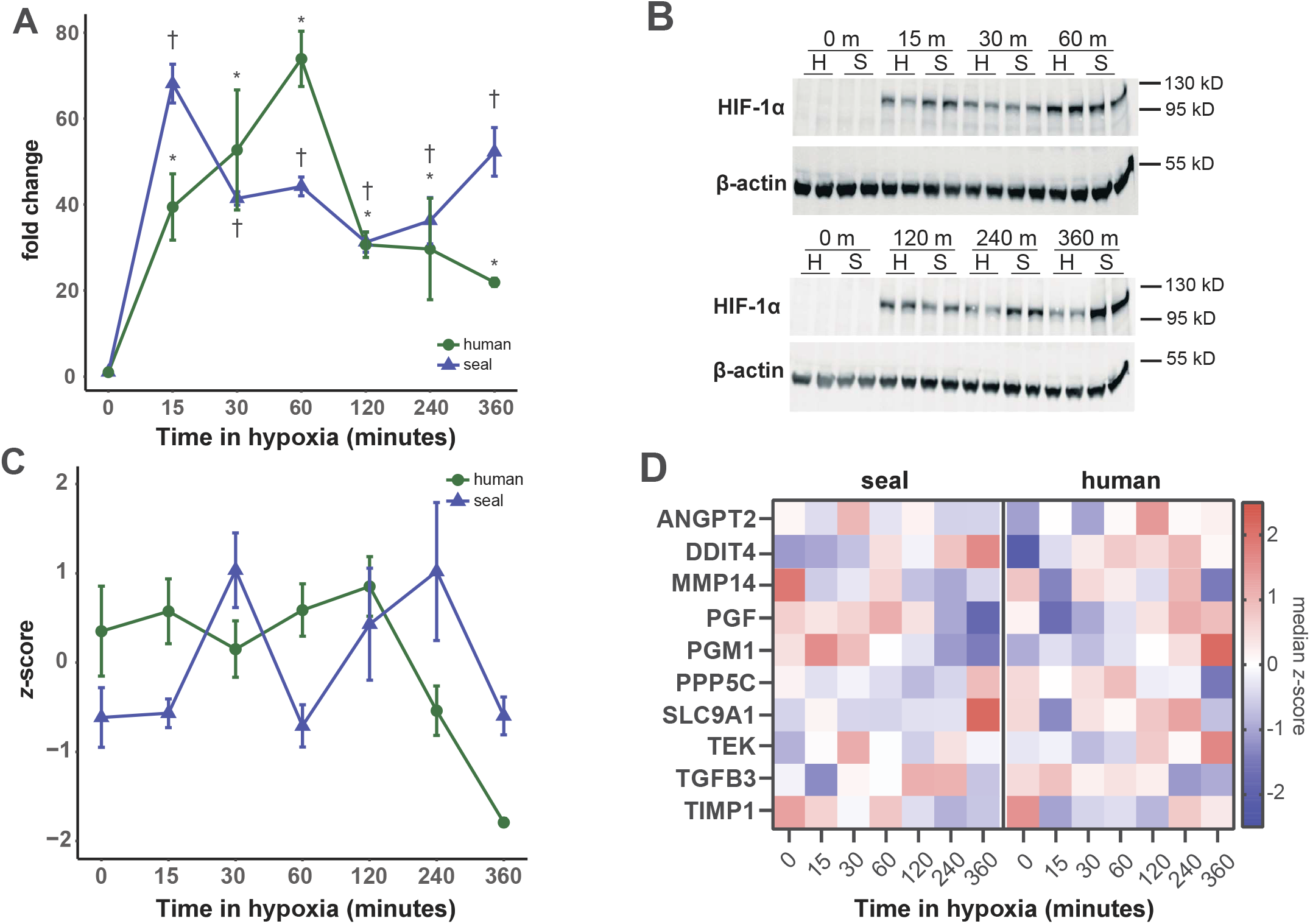
Hypoxia rapidly stabilizes HIF-1α in seal endothelial cells. (A) Fold change in HIF-1α protein abundance compared to normoxic baseline in endothelial cells exposed to 1% O_2_ for up to 6 h, n=3. † p<0.05 versus control (seal); * p<0.05 versus control (human). (B) Representative western blots showing HIF-1α abundance. H, human; S, seal. (C) *z-*score for HIF-1α mRNA levels in response to hypoxia exposure, n=3. (D) Median *z*-scores for select HIF-1 target genes in seal and human cells.

Seal and human cells displayed distinct gene expression patterns for several HIF-1 targets. Expression of factors implicated in extracellular matrix reorganization, angiogenesis and cell motility were elevated across the time course in human cells but were generally repressed or lagged behind in seal cells, including angiopoietin 2 (ANGPT2), the angiopoietin 1 receptor (TEK; TIE2), the Na+/H+ antiporter (SLC9A1; NHE1), protein phosphatase 5 catalytic subunit (PPP5C), transforming growth factor beta-3 (TGFB3), and matrix metalloprotease 14 (MMP14), while a metalloprotease inhibitor (TIMP1) showed the opposite expression pattern (Figure 2D). Placental growth factor (PGF) was downregulated in seal cells after 2 h hypoxia but displayed the opposite trend in human cells. Expression of the cellular stress protein DNA damage-inducible transcript 4 (DDIT4; REDD1) increased in seal cells at late timepoints, but its expression lagged that in human cells (Figure 2D). Seal cells increased expression of the first enzyme in glycogen synthesis, phosphoglucomutase 1 (PGM1), at early timepoints while it was generally repressed in human cells (Figure 2D). Together, these results suggest that the angiogenic signaling pathway is decoupled from HIF-1α stabilization in seal cells during hypoxia exposure.

### Seal cells delay the onset of pro-angiogenic transcriptional programs during hypoxia

We used RNAseq to evaluate global changes in gene expression in seal and human cells during a 6 h hypoxia exposure. We conducted *cis-*regulatory analyses with differentially expressed (DE) genes to predict transcription factors (TFs) that play major roles in the response to hypoxia at each time point. Comparative analyses of the top three predicted TFs in seal and human cells after 30 min, 60 min, and 6 h in hypoxia revealed species-specific dynamics in transcriptional control: none of the nine predicted seal TFs overlapped across time points, while only six distinct TFs were predicted in human cells (Figure 3A,B). Two TFs (chromodomain helicase DNA binding protein 1, CHD1 and serum response factor, SRF) were represented in both human and seal datasets. Predicted TFs in seal cells at 30 and 60 min suggest rapid metabolic (forkhead box K1, FOXK1) and inflammation-related (NF-κB subunit RELA; NFKB1; JUN; zinc finger and BTB containing 7A, ZBTB7A) responses to hypoxia, with a limited angiogenic signature (JUN). In contrast, the transcriptional signature in human cells at both 30 and 60 min suggests altered expression of oxygen delivery pathways via homeobox A13 (HOXA13), FOS like 1 (FOSL) and SRF, all of which regulate angiogenesis, vascular remodeling, and vasodilation (Evellin et al., 2013; Franco and Li, 2009; Galvagni et al., 2013; Shaut et al., 2008). In addition, broad changes in DNA replication and transcription may characterize the human cell hypoxia response via churchill domain containing 1 (CHURC1) and CHD1.

**Figure 3.**
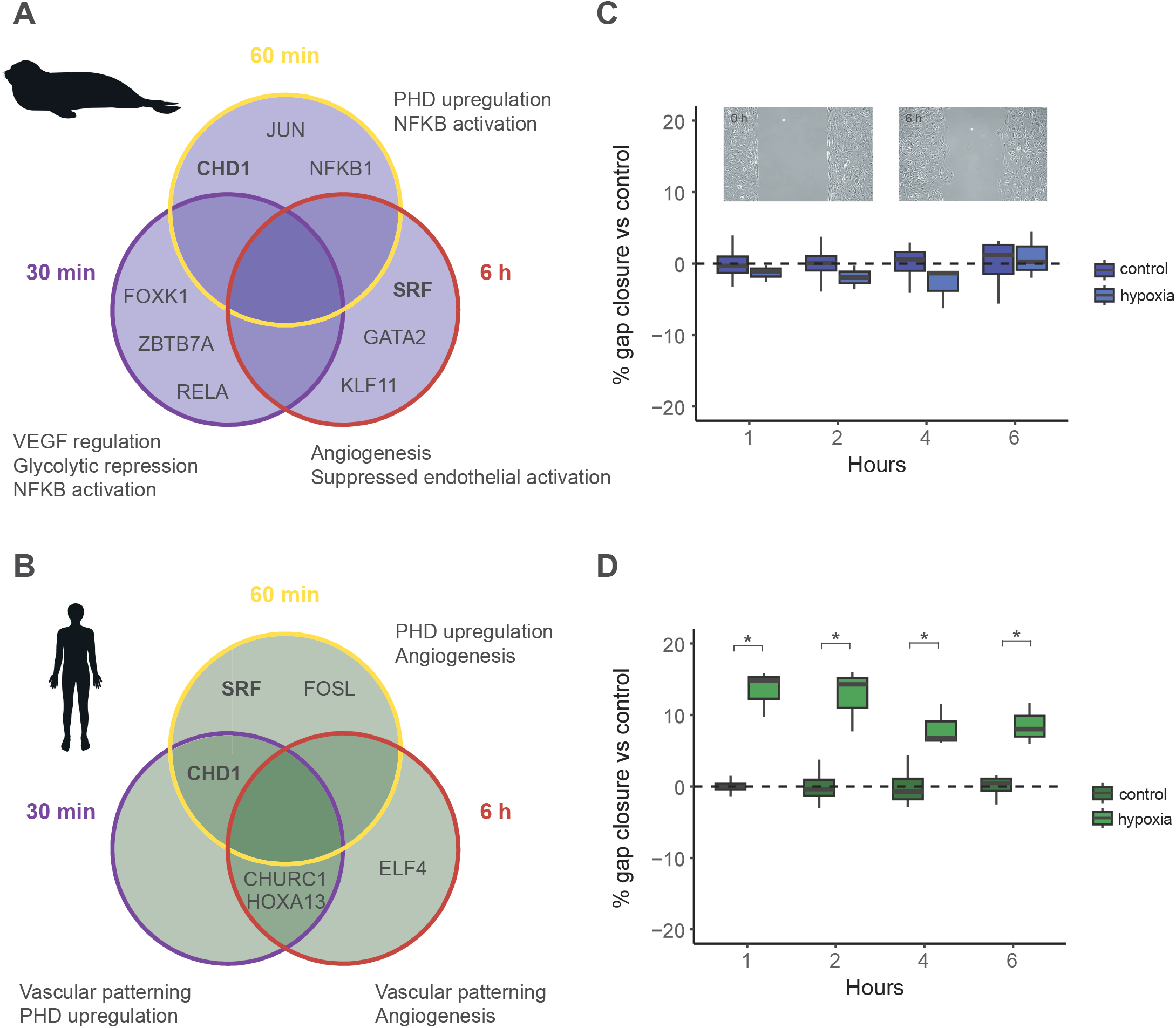
Seal cells delay angiogenic signaling and do not increase cell migration in response to hypoxia exposure. (A, B) Top three transcription factors predicted from genes DE at 30 min, 60 min, or 6 h versus control for (A) seal and (B) human cells. Bold text indicates factors shared between species. (C, D) Rates of gap closure in (C) seal and (D) human endothelial cells in hypoxia compared to control. Data are normalized to normoxic controls at respective timepoints to account for overall cell migration over time. Data are n=4 dishes from n=2 independent experiments. * p<0.05 vs control. Inset: representative images of scratch at 0 h and after 6 h cell migration. Scale bar is 50 µm.

Seal cells do appear to regulate expression of genes involved in oxygen delivery during hypoxia at 6 h. Two of the top three predicted TFs in seal cells at 6 h (GATA binding protein 2, GATA2; SRF) regulate angiogenesis, vascular remodeling, and vascular tone (Franco and Li, 2009; Yamashita et al., 2001). The third TF, Kruppel like factor 11 (KLF11) maintains vascular homeostasis and limits endothelial oxidant stress and inflammation via inhibition of NADPH oxidase 2 (NOX2) and matrix metalloproteinase 9 (MMP9) expression (Fan et al., 2013; Zhao et al., 2021). In human cells, repeated identification of HOXA13 and CHURC1 suggests a continued angiogenic signal at 6 h along with upregulation of transcription. Additionally, ELF4-mediated transcriptional changes may promote cell cycle entry during hypoxia exposure (Sivina et al., 2011; Suico et al., 2016). Cell migration data support these transcriptional data, as hypoxia accelerated migration into a cell-free gap in human but not seal cells over a 6 h exposure (Figure 3C,D). Together, these results suggest a highly dynamic transcriptional response to hypoxia in seal cells which alters cellular metabolism to promote endothelial homeostasis. This response stands in contrast to the consistently pro-angiogenic signal observed in human cells exposed to hypoxia.

### The transcriptional response to hypoxia highlights species-specific differences in antioxidant gene expression

Hypoxia rapidly repressed global gene expression in both seal and human cells, with 70% of differentially expressed (DE) genes in seal and 60% of DE genes in human cells downregulated at 15 min compared to control (p<0.001 for both species) (Figure 4). Expression in seal cells remained repressed from 30 min until 4 h (p<0.005 at each time point), at which point up-versus down-regulation did not differ from 50/50 (4 h: p=0.68; 6 h: p=0.36) (Figure 4A). In human cells, repression was similarly maintained through 1 h (p<0.001 at each time point), did not differ from 50/50 at 2 h (p=0.16), and shifted toward upregulation from 4-6 h (4 h: p<0.001; 6 h: p<0.005) (Figure 4C). In both species, DE gene expression decreased at 60 and 120 min (Figure 4B,D); in human cells this coincided with a shift toward upregulation (Figure 4C) while in seal cells gene expression patterns remained constant but with a tendency toward downregulation (Figure 4A).

**Figure 4.**
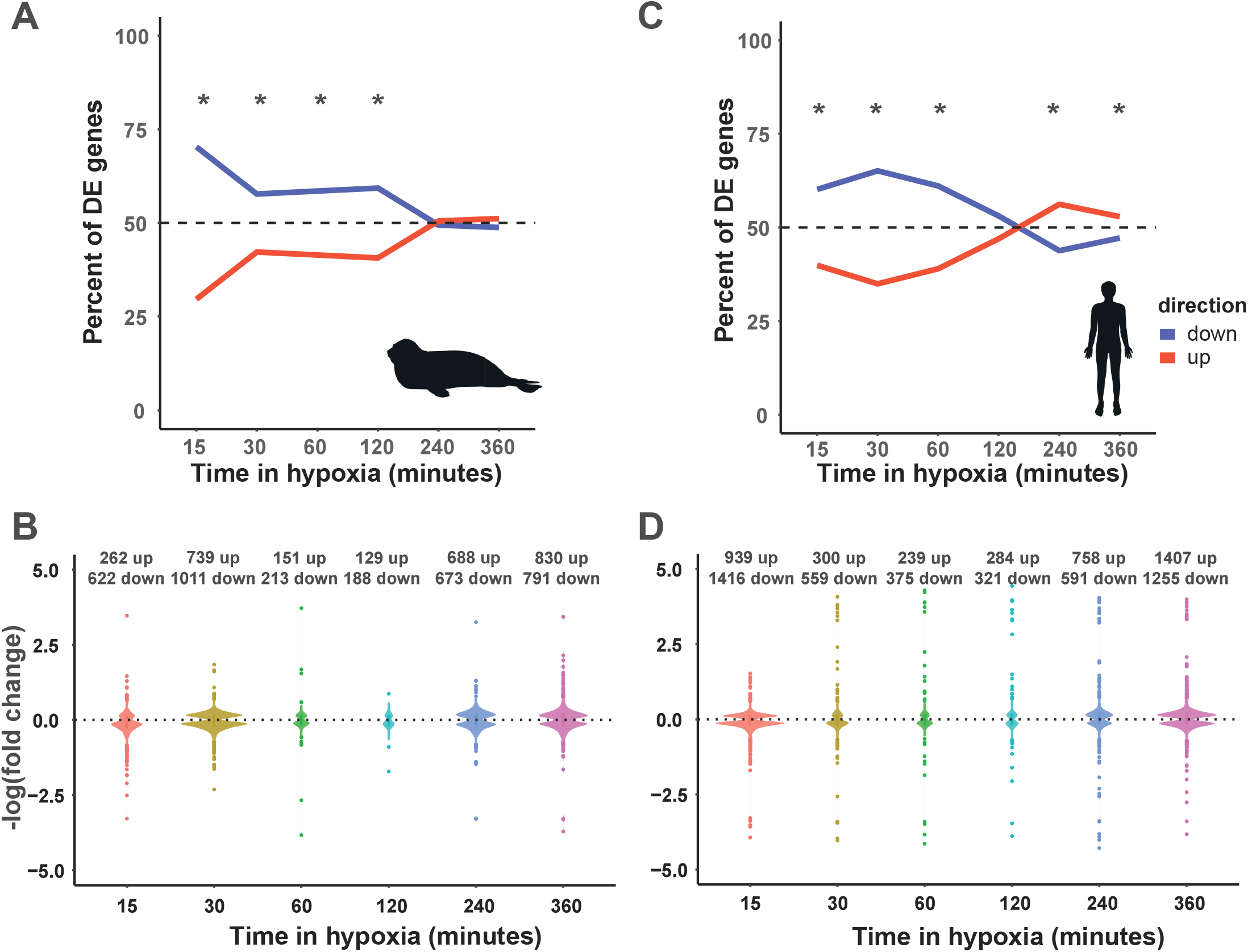
Seal cell gene expression equilibrates under sustained hypoxia exposure. Global differential gene expression patterns in seal (A, B) and human (C, D) cells during hypoxia compared to normoxic control. * p<0.05 compared to 50/50.

In both species, the least number of DE genes were detected at 60 and 120 minutes (Figure 4B,D); in human cells this coincided with a shift toward upregulation (Figure 4C) while in seal cells it overlapped with a plateau during which downregulation dominated (Figure 4A).

We then identified concerted changes in gene expression using *k*-means clustering (Figure 5A) and conducted GSEA for KEGG pathways within each cluster (Figure 5B; Figure S1). Seal cluster 6 and human cluster 3 were both enriched for HIF-1 signaling. Seal cluster 6 contained genes which gradually increased over time, while human cluster 3 genes exhibited a gradual increase in expression until 4 h, followed by a rapid decrease at 6 h. Seal cluster 5 and human cluster 6 were enriched for TNF signaling; both clusters displayed a rapid decrease in expression followed by a gradual return to baseline, suggesting that both human and seal cells downregulate pro-inflammatory pathways during the initial response to hypoxia.

**Figure 5.**
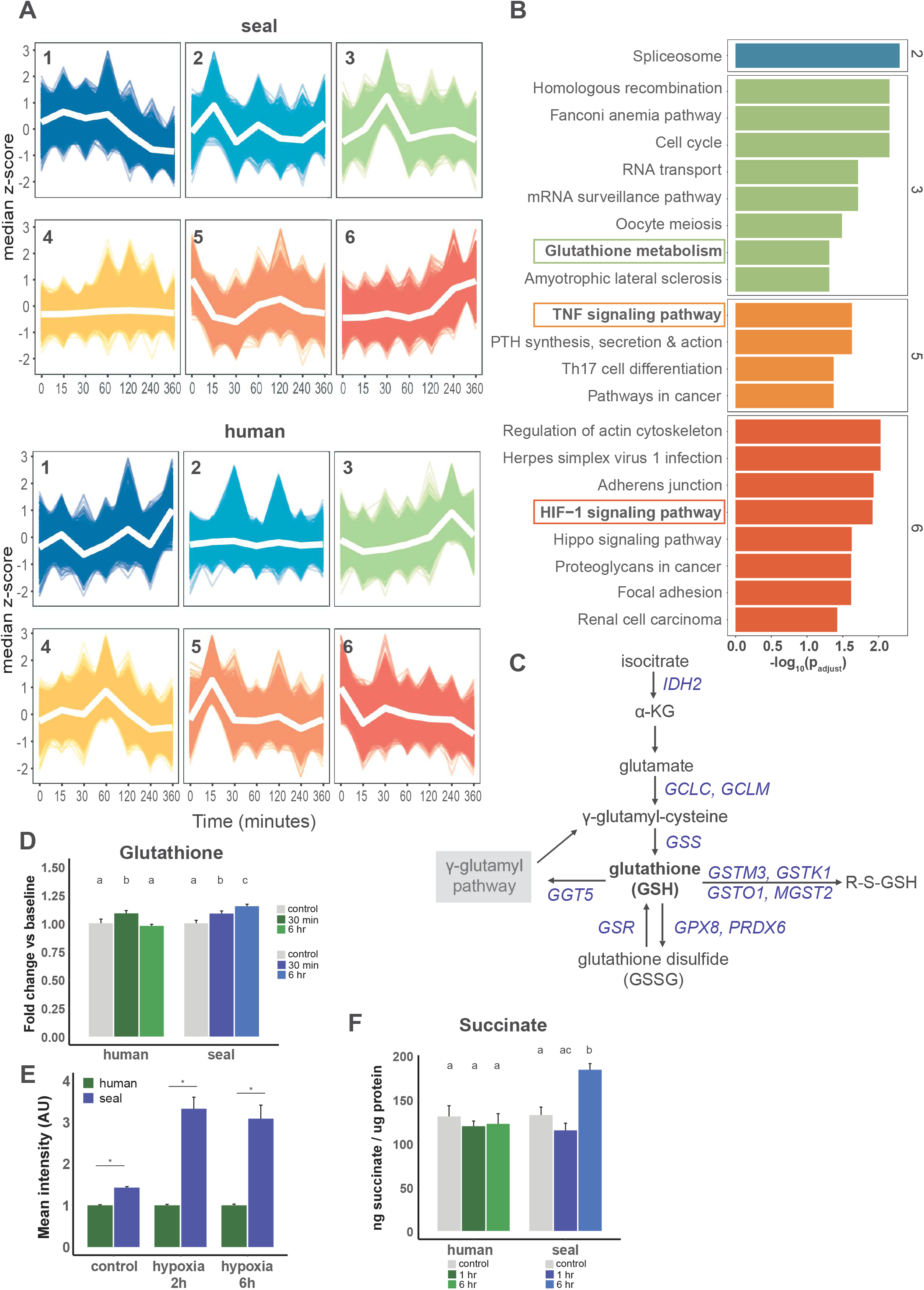
Concerted changes in gene expression in response to hypoxia in seal and human cells. (A) *k*-means clustering of gene expression data for seal (left) and human (right) cells exposed to 1% O_2_ for up to 6 h. (B) KEGG pathway enrichment for seal clusters. Numbers on the right axis correspond to seal cluster numbers. (C) Diagrammatic pathway of glutathione synthesis. Blue italicized gene symbols represent genes in seal cluster 3. See also Figure S1. (D) GSH content relative to each species’ baseline. Lettering indicates intraspecific changes among treatments. (E) Mean intensity of intracellular ThiolTracker Violet fluorescence during hypoxia exposure. Data are normalized to human at each timepoint. * p<0.05 between species. (F) Percent change in intracellular succinate concentration in response to hypoxia exposure. * p<0.05 versus species baseline.

Seal cluster 3 was enriched for glutathione (GSH) metabolism; this cluster contained genes for which expression peaked within 30 min of hypoxia onset. No human gene expression cluster displayed enrichment for genes involved in GSH metabolism (Figure S1). Genes from the GSH metabolism pathway present in seal cluster 3 are listed in Table 1 and include the critical GSH biosynthesis genes glutamate-cysteine ligase catalytic (GCLC) and modifier (GCLM) subunits and glutathione synthetase (GSS) (Figure 5C). GSH levels transiently increased in human cells during hypoxia exposure but continuously increased in seal cells through 6 h in hypoxia (human: F_2,20_=4.034, p=0.04; seal: F_2,21_=11.03, p=0.0005) (Figure 5D). Furthermore, baseline GSH content was ∼1.4-fold greater in seal compared to human cells (t=13.26, p<0.001) and remained ∼3-fold higher in seal than in human cells during hypoxia exposure (2 h: 3.3-fold, t=8.262, p=0.014; 6 h: 3.1-fold, t=6.422, p=0.022) (Figure 5E), suggesting that early transcriptional changes during hypoxia exposure support increased GSH pools in seal but not human cells.

**Table 1.** Glutathione metabolism genes in seal cluster 3.

| **Gene symbol** | **Gene name** |
| --- | --- |
| *IDH2* | isocitrate dehydrogenase (NADP(+)) 2 |
| *GCLC* | glutamate-cysteine ligase catalytic subunit |
| *GCLM* | glutamate-cysteine ligase modifier subunit |
| *GSS* | glutathione synthetase |
| *GGT5* | gamma-glutamyltransferase 5 |
| *GSR* | glutathione-disulfide reductase |
| *PGD* | phosphogluconate dehydrogenase |
| *PRDX6* | peroxiredoxin 6 |
| *GPX8* | glutathione peroxidase 8 |
| *GSTO1* | glutathione S-transferase omega 1 |
| *GSTM3* | glutathione S-transferase mu 3 |
| *GSTK1* | glutathione S-transferase kappa 1 |
| *MGST2* | microsomal glutathione S-transferase 2 |
| *SRM* | spermidine synthase |
| *SMS* | spermine synthase |
| *ODC1* | ornithine decarboxylase 1 |
| *RRM1* | ribonucleotide reductase catalytic subunit M1 |
| *RRM2B* | ribonucleotide reductase regulatory TP53 inducible subunit M2B |

Several genes involved in mitochondrial metabolism and redox homeostasis were also present in seal cluster 3, including ornithine decarboxylase 1 (ODC1) spermidine synthase (SRM), and spermine synthase (SMS) (Puleston et al., 2019; Qiu et al., 2022; Rigobello et al., 1993). Mitochondrial succinate accumulation during ischemia (and associated tissue hypoxia) drives production of superoxide via reverse electron transport at mitochondrial complex I upon reoxygenation (Chouchani et al., 2014; Chouchani et al., 2016), thus we measured intracellular succinate levels during hypoxia exposure. Interestingly, succinate levels increased 40% in seal but not human cells after 6 h hypoxia (seal: F_2,6_=19.30, p=0.0024; human: F_2,6_=0.3164, p=0.74) (Figure 5F); this relatively mild accumulation may support continued oxidative phosphorylation in seal cells after hypoxia/reoxygenation as observed in Figure 1D. Peroxiredoxin 6 (PRDX6), microsomal glutathione S-transferase 2 (MGST2), and gamma glutamyl transferase 5 (GGT5), which contribute to glutathione-dependent leukotriene synthesis (Miyata et al., 2020; Wickham et al., 2011) and may promote vasoconstriction in diving mammals (Blawas et al., 2021), were also present in seal cluster 3. Together, these data suggest that in contrast to human cells, seal endothelial cells maintain cellular GSH levels, remain metabolically active, and maintain vasoconstrictive signaling during hypoxia exposure.

### The short-term transcriptional response to hypoxia in seal cells limits inflammatory signaling

We next investigated whether short-term hypoxia (≤ 1 h) leads to differential functional enrichment in gene expression between species. We detected 42 DE genes shared between the seal and human cell response to short-term hypoxia. Generally, these shared genes displayed similar expression patterns between species (Figure 6A). Seal genes responsive to short-term hypoxia were enriched for several TGF-β signaling pathways including downregulation of TGF-β receptor signaling. In contrast, the response in human genes was dominated by pathways involved in RNA processing and transcription, ER stress, and upregulation of electron transport chain subunits (Figure 6B, Table S2). Functional interaction networks (FINs) suggested a central role for NFKB1 (p50) in the short-term hypoxia response in both species, though expression of this subunit and its related RELB subunit were downregulated relative to control in both species (Figure 6CD). The seal FIN also confirmed a critical role for downregulation of TGF-β signaling in seal cells (Figure 6C) as the only upregulated gene in the network was MTMR4, a TGF-β inhibitor. Interestingly, the angiogenic gene expression signature in seal cells suggested blunting of VEGF-dependent angiogenesis during short-term hypoxia (Figure 6C).

**Figure 6.**
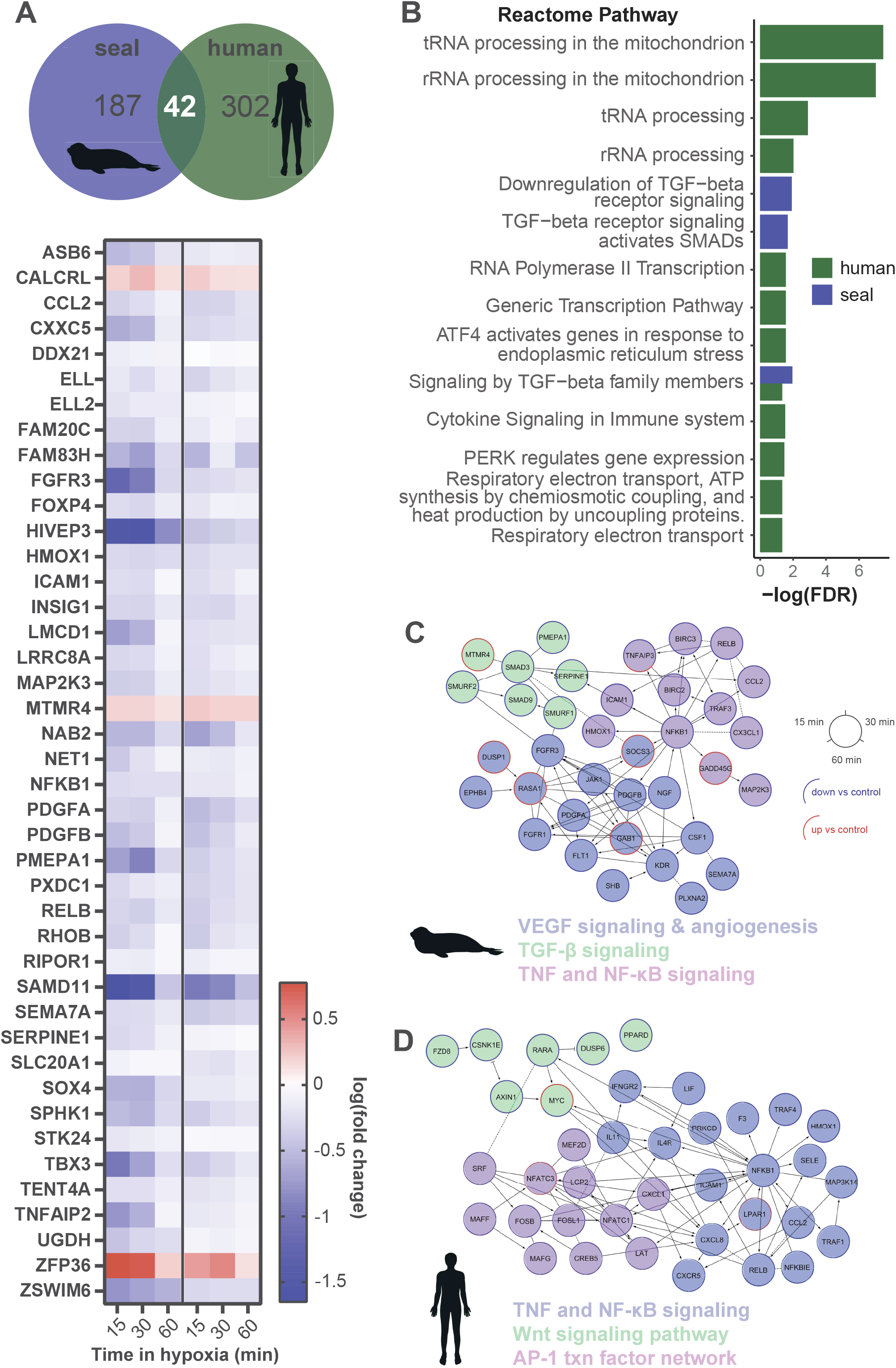
Short-term hypoxia modulates inflammatory signaling in seal cells. (A) Overlap between genes DE at all early time points (15 min, 30 min 60 min) versus control for seal and human cells. Heatmap log_10_(fold change) for 42 DE genes shared between species. (B) Reactome pathway enrichment for genes DE at all early time points. (C,D) Functional interaction networks and select enriched pathways for seal (C) and human (D) genes DE at all early time points.

### Pro-proliferative transcriptional signals characterize the long-term hypoxia response in seal cells

We next evaluated expression patterns in response to hypoxia exposure for ≥ 2 h. DE genes shared between seal and human cells generally displayed similar trajectories, apart from HOXA13 (homeobox A13; also predicted as coordinating the human response to hypoxia in cells (Figure 3C)) for which long-term hypoxia repressed expression in seal but stimulated expression in human cells (Figure 7A). No Reactome pathways were overrepresented in the 193 seal genes DE for 2-6 h hypoxia exposure. Human DE gene expression (333 genes) was again enriched for pathways related to tRNA and rRNA processing and respiratory electron transport (universally upregulated as during short-term exposure) as well as negative regulation of the MAPK pathway (Figure S2A; Table S3).

**Figure 7.**
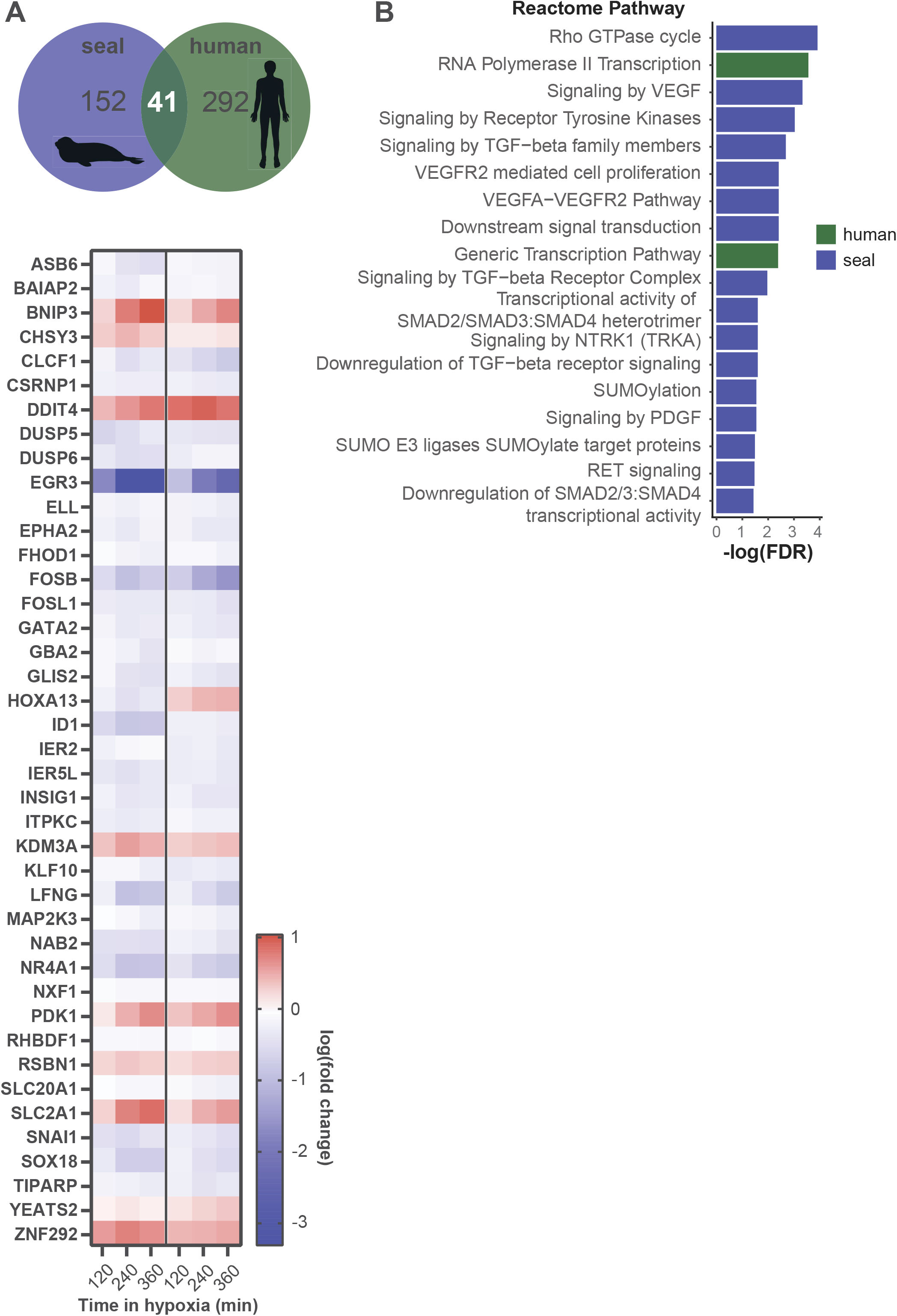
Long-term hypoxia modulates cell proliferation pathway expression in seal cells. (A) Overlap between genes DE at all late time points (2 h, 4 h, 6 h) versus control for seal and human cells. Heatmap shows log_10_(fold change) for 41 DE genes shared between species. (B) Reactome pathway enrichments for genes DE at 6 h versus control.

We then separately considered the transcriptional response to 6 h hypoxia, at which point a second increase in seal HIF-1α protein abundance occurred alongside species-specific expression patterns for several HIF-1 targets (Figure 2A,D). GSEA revealed downregulation of several pathways related to transcription and mRNA splicing, as well as C-type lectin receptors (implicated in the immune response, cell adhesion and apoptosis; (Chiffoleau, 2018)) in human cells at 6 h (Figure S2B). Reactome pathway enrichment analyses further supported hypoxia-induced changes in transcription including RNA polymerase II activity in human cells (Figure 7B). In contrast, seal cell gene expression at 6 h was enriched for several Smad and TGF-β signaling pathways, the VEGFA-VEGFR2 pathway, SUMOylation and SUMO E3 ligase target proteins (Figure 7B). Hypoxia-induced suppression of TGF-β signaling appeared to be regulated through increased expression of the Smad phosphatase MTMR4 and the ubiquitin ligase NEDD4L, which targets TGF-β-induced pSmad2/3 (Table S4) (Gao et al., 2009; Yu et al., 2010). Despite the observed shift toward pro-proliferative signaling during long-term hypoxia exposure, we did not detect changes in cell migration in seal endothelial cells in response to hypoxia (Figure 3C,D).

## Discussion

Elephant seals repeatedly experience profound hypoxemia during continuous diving (Meir et al., 2009; Meir et al., 2013). While similar fluctuations characterize pathophysiological events in non-hypoxia adapted species (Eltzschig and Eckle, 2011; Kalogeris et al., 2012), the cellular and molecular mechanisms which confer hypoxia tolerance in seals remain unclear. Here, we developed a proliferative arterial endothelial cell culture system using cells derived from elephant seals to identify key transcriptional regulators of the response to hypoxia. Our results show blunted angiogenic signaling despite rapid and sustained HIF-1α stabilization in seal cells. Moreover, seal cells also increase GSH pools during hypoxia, which may mitigate oxidant stress upon reoxygenation despite requiring energetic input under oxygen-limited conditions. High maximal and spare respiratory capacity at baseline and after hypoxia/reoxygenation in seal cells may support such demands. Together, these data suggest that seal endothelial cells modulate metabolic parameters to meet oxygen availability during hypoxia rather than attempting to increase oxygen delivery in the face of limited supply.

### Canonical hypoxia signaling is decoupled from angiogenesis in seal cells

HIF-1 is the “master regulator” of the canonical response to hypoxia (Pugh and Ratcliffe, 2003; Wang et al., 1995), typically promoting both angiogenesis and glycolysis. *In vivo, in vitro*, and *in silico* analyses of HIF-1α structure and function indicate that marine mammal HIF-1α stabilization is highly sensitive to hypoxia (Bi et al., 2015; Johnson et al., 2004; Johnson et al., 2005; Penso-Dolfin et al., 2020; Vázquez-Medina et al., 2011c). HIF-1α stabilization occurred rapidly in seal endothelial cells exposed to hypoxia, and high levels of the protein persisted in seal cells through a 6 h exposure. Interestingly, seal cell expression of several HIF-1 targets implicated in vascular homeostasis (ANGPT2, MMP14, SLC9A1) (Cork et al., 2012; Yang and Rosenberg, 2015; Yu et al., 2008; Yuan et al., 2021) remained stable during long-term hypoxia exposure in seal cells, while expression was variable or increased in human cells. Global transcriptional patterns supported this regulation as DE genes in seal cells predicted factors which regulate cell proliferation (FOXK1, ZBTB7A) and suppress endothelial activation (KLF11) (Fan et al., 2013; Yin et al., 2013), while a human cells present a consistently pro-angiogenic signal despite slower HIF-1α stabilization. Accordingly, hypoxia increased gap closure rates in human but not seal cells. These results suggest that seal cells rapidly stabilize HIF-1α but further regulate expression of putative HIF-1 targets during hypoxia exposure.

### Seal cells dampen inflammatory signaling throughout hypoxia exposure

Our data suggest continued regulation of inflammatory signaling in seal endothelial cells across 6 h of hypoxia exposure. Gene expression signatures early (≤ 1 h) in hypoxia exposure highlight downregulation of the TNF, NF-ĸB, and TGF-β pathways in seal cells. In contrast, enrichments on human cell gene expression identified substantially fewer inflammatory pathways. Continued overrepresentation of TGF-β-related signaling pathways (including SMAD activity) coupled with increased expression of negative regulators including the dual specificity protein phosphatase MTMR4 and ubiquitin ligase NEDD4L in seal cells at 6 h, along with ZBTB7A at 30 min suggests that tight regulation of the TGF-β signaling cascade is critical in the seal cell response to hypoxia. Our data also suggest that altered NF-ĸB signaling during hypoxia exposure in seal cells may be regulated by TNF. NF-ĸB activation may be either pro-inflammatory or pro-survival depending on cellular context. Activity of both NF-ĸB and its inhibitor IKKβ are required for hypoxic HIF-1α accumulation and HIF-1 target gene expression (Rius et al., 2008). Knockdown of the NF-ĸB subunit RelA in endothelial cells impairs smooth muscle cell proliferation (Patel et al., 2017) and NF-ĸB activity is implicated in eNOS downregulation during chronic intermittent hypoxia (Wang et al., 2013). In Weddell seals, nitric oxide sensitivity varies across tissues and vessel beds but is generally blunted when compared to sheep (Hindle et al., 2019), presumably to avoid hypoxic vasodilation during diving. Here, we found that tight regulation of NF-ĸB expression in seal cells during hypoxia exposure may promote cell survival and modulate vasodilatory signaling in the endothelium. These results corroborate a generally reduced inflammatory response in whole blood from elephant seals stimulated with lipopolysaccharide (Bagchi et al., 2018). In contrast, changes to inflammatory signaling pathways occurring in human cells during short-term hypoxia exposure are supplanted by broad changes in transcriptional regulation during long-term exposure.

### Hypoxia promotes GSH synthesis in seal cells

High tissue and circulating GSH levels, as well as high activity of many glutathione-dependent antioxidant enzymes, including glutathione peroxidases (GPX) and glutathione *S*-transferases (GST) (Cantú-Medellín et al., 2011; Murphy and Hochachka, 1981; Vázquez-Medina et al., 2007; Vázquez-Medina et al., 2011a; Vázquez-Medina et al., 2011b), suggest a critical role for GSH in protecting diving mammals against oxidative stress. Recent genomic work demonstrates selection on several genes in the glutathione metabolism pathway in marine mammals, including those we identified as enriched (Foote et al., 2015; Tian et al., 2019; Tian et al., 2021; Yim et al., 2014). To date, however, the only measurement of diving GSH levels is in Weddell seals, with the observation that near depletion of circulating GSH at the halfway point of restrained dives was followed by an increase above baseline levels within ∼5 min of recovery (Murphy and Hochachka, 1981). Interestingly, we found that elephant seal vascular endothelial cells continue GSH synthesis during long-term hypoxia exposure alongside rapid expression increases in several GSH biosynthesis and antioxidant enzymes. Hypoxic GSH accumulation in seal cells likely counteracts oxidant generation upon reoxygenation, protecting cells against oxidative damage and cellular stress. These results track with existing *in vivo, in vitro*, and *in silico* evidence supporting GSH’s role in maintaining redox homeostasis during diving in marine mammals (Cantú-Medellín et al., 2011; Murphy and Hochachka, 1981; Tian et al., 2019; Tian et al., 2021; Vázquez-Medina et al., 2007; Vázquez-Medina et al., 2011a; Vázquez-Medina et al., 2011b). Additionally, data from non-diving hypoxia tolerant species indicate that increases in GSH prior to oxidative insult protect against later tissue injury (Buzadžić et al., 1990; Hermes-Lima and Storey, 1993), potentially justifying the “cost” of producing GSH in oxygen-limited conditions.

### Hypoxic succinate accumulation may promote mitochondrial function in seal cells

In addition to GSH biosynthesis, other components of the glutathione metabolism pathway enriched in seal cells included those related to polyamine synthesis (via ODC1, SRM, and SMS) which may further protect cells against oxidant generation upon reoxygenation (Ha et al., 1998; Rider et al., 2007). Polyamines regulate the mitochondrial permeability transition pore *via* calcium flux, thus modulating activity of the pyruvate dehydrogenase complex and impacting mitochondrial respiration (Elustondo et al., 2015). Hypoxic accumulation of succinate in seal cells may derive from conversion of the polyamine putrescine to succinate and sustain oxidative phosphorylation during hypoxia exposure. While excessive succinate accumulation due to reversal of succinate dehydrogenase (SDH) activity during hypoxia drives superoxide production *via* complex I (Chouchani et al., 2016), inhibition of SDH activity by itaconate may promote mild succinate accumulation without associated superoxide generation (Cordes et al., 2020). Additionally, succinate competitively inhibits prolyl hydroxylases such as those responsible for HIF-1α hydroxylation (Chakrabarty and Chandel, 2022), supporting increased HIF-1α sensitivity to hypoxia in seal cells. Early increases in the HIF-1 target phosphoglucomutase 1 (PGM1) in seal cells may regulate glucose metabolism during hypoxia, consistent with glycolytic repression *via* ZBTB7A transcriptional control (Bae et al., 2014; Liu et al., 2014). Human cells, in contrast, upregulate several mitochondrial electron transport chain subunit components during short-term hypoxia exposure, consistent with increased mitochondrial branch length after 1 h, but overall changes in mitochondrial respiration in human cells following hypoxia/reoxygenation are milder than those in seal cells and are accompanied by higher ECAR, suggesting that human but not seal cells rely on glycolysis to support continued respiration during hypoxia.

## Conclusion

In summary, we found that seal endothelial cells tightly regulated inflammation and mitochondrial dynamics and blunted angiogenic signaling during hypoxia exposure. These changes occurred in parallel with increased GSH synthesis, which may protect seal cells against oxidant generation during reoxygenation. Conversely, human cells signaled for angiogenesis and vascular remodeling early and consistently throughout the exposure; these signals may destabilize the vasculature and increase susceptibility to inflammatory and oxidative damage to tissue during reoxygenation. Here we identify candidate pathways supporting endothelial cell hypoxia tolerance in a deep diving seal. Further investigation *via* co-culture with vascular smooth muscle, peripheral blood mononuclear cells, or serum from diving animals is required to anchor these responses in the context of the sustained vasoconstriction characteristic of the diving response.

## Acknowledgements

We thank Dr. Fung Lam for his assistance in obtaining human placental tissue. We thank Drs. Daniel Costa, Patrick Robinson, and Rachel Holser for collecting and sharing elephant seal placental tissue. We thank Dr. Rohit Kolora for improving the elephant seal genome annotation. The UC Berkeley QB3 Genomics facility provided sequencing service and support (RRID:SCR_022170; NIH S10 OD018174 Instrumentation Grant). Confocal imaging was conducted using resources at the UC Berkeley Molecular Imaging Center (RRID:SCR_017852), supported by the Gordon and Betty Moore Foundation.

## Methods

### Tissue acquisition

Northern elephant seal *(Mirounga angustirostris)* placentae were obtained under NMFS permit # 19108. Whole placentae were collected after live births (< 1 h) from the beach at Año Nuevo State Park (San Mateo, CA) during the pupping season. The generation of cell lines was conducted under NMFS permit # 22479. De-identified human placentae were donated following uncomplicated live births at Sutter Health’s California Pacific Medical Center. All placentae were maintained at 4°C until time of dissection. All dissections were performed within 6 hours of collection.

### Primary cell isolation and culture

Arteries were dissected from placental tissue and rinsed with ice-cold sterile Hanks’ Balanced Salt Solution (HBSS; Gibco, Waltham, MA, USA). Arteries were cut open lengthwise, flushed with HBSS, and placed in a collagenase type 2 solution (500 U/mL in DMEM; Worthington Biochemical, Lakewood, NJ, USA) for 30 minutes in a tissue culture incubator (37°C, 5% CO_2_, humidified). The collagenase solution was blocked with complete growth medium. The endothelial surface of the arteries was scraped into tissue culture-treated dishes containing complete growth medium and transferred to a tissue culture incubator. Endothelial cell islands were visible in the culture dishes after 24 h. Growth medium was changed daily for the first 4-5 days at passage 0. Seal cells were maintained in DMEM (Gibco, catalog # 11885-084) supplemented with 10% fetal bovine serum (Seradigm, Avantor, Mexico), 10 mM HEPES (Gibco), 1% (1X) Antibiotic-Antimycotic (Gibco), and 4 ug/mL endothelial cell growth supplement (ECGS; Corning, Corning, NY, USA). Human cells were grown in commercial endothelial cell medium (Sciencell Research Laboratories, catalog # 1001, Carlsbad, CA, USA). Confluent cultures were cryopreserved at passage 1. Pooled stock cultures were created for human (n=2-3) and seal (n=3) cells. Northern elephant seal trophoblasts were isolated by mincing placental labyrinth in a collagenase type II solution, followed by 5 minutes in a tissue culture incubator. The collagenase solution was blocked with complete medium and passed through a 100 µm filter. Filtrate was centrifuged at 200 x *g* for 5 minutes at 4°C. The pellet was washed once in HBSS and resuspended in trophoblast growth medium (Sciencell Research Laboratories, catalog # 7121), then plated on tissue culture-treated dishes coated with collagen IV. All experiments were conducted with cells between passages 4 and 7. All cells were switched to the same medium formulation the day prior to experiments.

### Cell characterization

The endothelial phenotype of the preparations was confirmed by Dil-acetylated LDL uptake (Invitrogen, catalog # L3484), RT-qPCR for vascular endothelial cadherin (CD144) and platelet endothelial cell adhesion molecule-1 (PECAM-1), and immunostaining for PECAM-1 (CD31; Novus Biologicals, catalog # NB100-2284, 1:100). The Dil-acetylated LDL uptake assay was conducted following manufacturer instructions. RT-qPCR was conducted using our previously published methods (Torres-Velarde et al., 2021) with minor modifications in the reaction conditions: 1 min at 95°C followed by 40 cycles of 20 sec at 95°C, 30 sec at 60°C. Primer sequences are listed in Table S1. Immunostaining was conducted following the protocol described in Vázquez-Medina et al. (2016). Tube formation was assessed by plating 10,000 cells per well in a µ-slide 3D angiogenesis glass bottom slide (Ibidi, Fitchburg, WI, USA, catalog # 81506) coated with Matrigel. All imaging was conducted using a Zeiss Axio Observer 7 inverted microscope fitted with 10X and 20X objectives.

### Extracellular flux assays

Oxygen consumption was measured using the Seahorse Mitochondrial Stress Test Kit (Agilent Technologies, Santa Clara, CA, USA) and an XFp Extracellular Flux Analyzer. 30,000 cells/well were seeded in fibronectin-coated miniplates. Cells were washed with serum-free assay medium (Seahorse XF DMEM pH 7.4, 5.56 mM glucose, 1 mM pyruvate, 4 mM L-glutamine) and incubated at 37°C in a non-CO_2_ incubator for 1 h prior to assay. Oxygen consumption rates (OCR) were measured in the presence of oligomycin (1 µM), carbonyl cyanide-*p*-trifluoromethoxyphenylhydrazone (FCCP; 1 µM for human cells, 2 µM for seal cells), and rotenone/antimycin A (0.5 µM). Optimal FCCP concentrations were determined empirically for each species. Protein content was measured in each well at the end of the assay using the Qubit Protein Assay Kit (Molecular Probes, Eugene, OR, USA). Oxygen consumption rate (OCR) and extracellular acidification rate (ECAR) were normalized to protein content. Mitochondrial function was calculated according to Divakaruni et al. (2014) and normalized to basal respiration per well.

### Hypoxia exposure

Cells were incubated in an InVivO_2_ physiological cell culture workstation (Baker Ruskinn, Sanford, Maine, USA) under the following conditions: 1% O_2_, 5% CO_2_, 37°C for 15 min, 30 min, 1 h, 2 h, 4 h and 6 h. Protein and RNA were collected within the hypoxic environment. Control cells remained in a standard tissue culture incubator (21% O_2_, 37°C, 5% CO_2_). Cell viability was confirmed in cells undergoing hypoxia exposure using a commercial viability/cytotoxicity kit (Thermo Fisher, catalog number: R37601). In some experiments, extracellular flux assays were conducted after hypoxia/reoxygenation treatments consisting of 1 h at 0.5% O_2_ (37°C, 5% CO_2_) followed by 30 min in room air (37°C, no CO_2_) prior to assay.

### Immunoblotting

Western blot was conducted using our previously published methods (Vázquez-Medina et al., 2019). Briefly, cells were scraped in DPBS (Gibco) containing 1% Triton X-100 and 2% Halt Protease and Phosphatase Inhibitor Cocktail (Thermo Scientific, Waltham, MA, USA). Lysates were sonicated and centrifuged. Total protein content in the supernatant was determined using a BCA Rapid Gold Protein Assay (Pierce, Rockford, IL, USA). Proteins were resolved using SDS-PAGE and transferred onto nitrocellulose membranes. Membranes were blocked with Odyssey blocking buffer (LICOR, Omaha, NE, USA) and incubated overnight with an antibody against hypoxia inducible factor-1α (HIF-1α, Cell Signaling Technology #36169; 1:500). Proteins were visualized using IRDye 800CW secondary antibodies (LICOR) and a two-color near-infrared system (Azure c500, Azure Biosystems, Dublin, CA, USA). Membranes were stripped and reprobed with an antibody against β-actin (Cell Signaling Technology #4967; 1:2,000). Intensity of individual bands was quantified using FIJI v2.1.0 and normalized to β-actin.

### Biochemical assays

Glutathione content was measured under normoxic conditions and after exposure to 30 min and 6 h hypoxia using the GSH-Glo Assay Kit (Promega, Madison, WI, USA) according to manufacturer instructions. Luminescence was measured using a SpectraMax M3 Microplate Reader (Molecular Devices, San Jose, CA, USA). Changes in GSH levels were also measured in hypoxic cells using ThiolTracker Violet (Invitrogen, catalog # T10096). Cells were loaded with 20 µM ThiolTracker Violet glutathione detection reagent and 100 nM SYTO 16 Green Fluorescent Nucleic Acid Stain (Invitrogen, catalog # S7578) and imaged within the hypoxic environment using an LS620 microscope (Etaluma, Carlsbad, CA, USA) fitted with a 20X objective at 0, 30 minutes, and 6 hours at 1% O_2_. Fluorescence intensity was quantified in 12 random cells per field using FIJI v2.1.0.

Intracellular succinate concentration was measured in cells under normoxic conditions and after exposure to 1 and 6 h hypoxia using a commercial colorimetric assay (Sigma catalog # MAK184, St. Louis, MO, USA) according to manufacturer instructions. Absorbance at 450 nm was measured using a SpectraMax M3 Microplate Reader (Molecular Devices, San Jose, CA, USA). Succinate concentrations per well were normalized to protein content using a Pierce Rapid Gold Protein Assay (Pierce, Rockford, IL, USA).

### Cell migration assay

Cells were plated in an Ibidi 2-well silicone culture insert (Ibidi, catalog # 81176) adhered to a TC-treated 35 mm glass bottom dish and grown to confluence. At confluence, culture inserts were removed to create a “scratch.” Dishes were washed with DPBS to remove unattached cells, and media was changed. Dishes were incubated in control (normoxia) or hypoxia (1% O_2_) conditions for 6 h. Live cells were imaged using a Zeiss Axio Observer 7 inverted microscope fitted with a 10X objective at 1, 2, 4, and 6 h. Cell migration was calculated as the percent difference in cell-free area in hypoxic dishes compared to control at each time point using the ScratchAssayAnalyzer function in the MiToBo FIJI plugin.

### Mitochondrial staining and network analysis

Live cells were stained with MitoTracker Red CMXRos (Cell Signaling Technology, Danvers, MA) and NucBlue (Hoescht 33342, Invitrogen) under normoxic conditions and after 1 and 6 h hypoxia (1% O_2_) followed by 30 min reoxygenation. Cells were fixed in ice cold 1:1 methanol:acetone, rinsed with PBS, and imaged in VECTASHIELD mounting media (Vector Laboratories, Burlingame, CA) using a Zeiss LSM 710 microscope fitted with a 40X objective (Plan-Apochromat 1.4 NA oil, 0.13 mm). Mitochondrial footprint and mean branch length were determined from unfiltered binarized images using the MiNA (v2.0) FIJI plugin (https://github.com/StuartLab/MiNA).

### RNAseq

Hypoxic cells were lysed in Buffer RLT without β-mercaptoethanol (Qiagen, Germantown, MD, USA). RNA extractions were carried out using an RNeasy Mini Kit (Qiagen) including on-column DNase treatment according to manufacturer instructions. DNase treatment was confirmed by a lack of genomic amplification using PCR. RNA yield was quantified using a Nanodrop 1000 Spectrophotometer (Thermo Scientific). RNA integrity number (RIN) was determined using an Agilent 2100 Bioanalyzer kit for total eukaryotic RNA (Agilent Technologies, Santa Clara, CA, USA). Samples underwent library preparation and sequencing at the UC Berkeley Functional Genomics and Vincent J. Coates Genomics Sequencing Laboratories. A KAPA RNA HyperPrep Kit (catalog # KK8541; Roche, Basel, Switzerland) was used to prepare cDNA libraries from poly(A)-captured mRNA with TruSeq adapters. Three replicates per condition were used to generate libraries, with a sequencing depth of 25 M reads per sample on a NovaSeq platform (Illumina, San Diego, CA, USA).

### Transcriptome analyses

The elephant seal *(Mirounga angustirostris)* genome was annotated as described in our previous work (https://www.dnazoo.org/assemblies/Mirounga_angustirostris) (Torres-Velarde et al., 2021). Reads were mapped to the elephant seal or human genomes (version GRCh38) using STAR (Dobin et al., 2013).Transcript levels were quantified with RSEM v1.3.1 (Li and Dewey, 2011). Expression levels were converted to *z*-scores and normalized by the mean and variance across all replicates and conditions for each gene. The median *z*-score was selected for each gene and used to construct a Euclidean distance matrix. Genes that changed in a concerted manner across hypoxia exposure times were clustered by *k*-means. Optimal clustering was selected at *k*=6 due to diminishing returns at higher *k*. Within each cluster, gene set enrichment analysis (GSEA) with KEGG terms was used to identify enriched biological pathways. Additionally, genes differentially expressed between control and each timepoint were identified using EBSeq with a false discovery rate of 5% (Leng et al., 2013). GSEA for Reactome pathways was conducted using a summary list of genes differentially expressed at 15 min, 30 min and 1 h versus control (Subramanian et al., 2005). Functional Interaction Networks (FIN) were built for the same gene lists using Cytoscape v3.8.2 (Wu et al., 2017; Wu et al., 2010). *Cis*-regulatory analyses were conducted using iRegulon in Cytoscape (Janky et al., 2014).

### Statistical analyses

Statistical analyses were performed using GraphPad Prism v9.3.1. Normality was evaluated using D’Agostino-Pearson or Shapiro-Wilk tests, depending on sample size. When normality assumptions were met, differences between groups with equal variance were assessed by independent *t-*test or one-way ANOVA. When homoscedasticity assumptions were not met, differences between groups were determined by *t-*test with Welch’s correction or Welch’s ANOVA. In cases where data were not normally distributed, Kruskal-Wallis tests were used. In all cases, we corrected for multiple comparisons by setting FDR to 0.05; post hoc analyses used the two-stage linear step-up procedure of Benjamini, Krieger and Yekutieli. Data are presented as mean ± s.e.m.

---

**Figure S1. KEGG pathway enrichment for human clusters.** Numbers on the right axis correspond to human cluster numbers.

**Figure S2. Long-term hypoxia exposure modulates transcription and translation in human cells.** (A) Reactome pathway enrichments for genes DE at all late time points in human cells; no pathways were enriched in seal. (B) GSEA for genes DE at 6 h versus control in human cells; no pathways were enriched in seal. Normalized enriched score < 0 indicates net downregulation of the pathway.

