## Supplementary material for "Hypoxia blunts angiogenic signaling and upregulates the antioxidant system in elephant seal endothelial cells": Table S1

**Supplementary Table 1.** Primer sequences

| Gene | Sequence 5’ to 3’ |
|---|---|
| *ve-cadherin (cd144)* | CCC AGA ACC GGA TGA CCA AG |
| | TTT CGG ATG GAG ACG CTG CT |
| *pecam-1 (cd31)* | CAA TAG AAG GCG GGG TCG TG |
| | CGT GGC TTG GCA CTG GAA GT |
| *actin* | CGG TCA GTT CAT GGC TGA GG |
| | AAG GCT CGG ACC TTC CCA AC |
| *gapdh* | CAA GGC TGA GAA CGG GAA GC |
| | ATC GGC AGA GGG AGC AGA GA |
