## Supplementary material for "Hypoxia blunts angiogenic signaling and upregulates the antioxidant system in elephant seal endothelial cells": Table S2

**Table S2. Fold change in expression of respiratory electron chain components in response to short-term hypoxia exposure in human cells.** Fold change is expressed as a comparison to species baseline.

| Gene name | 15 min | 30 min | 60 min |
|---|---|---|---|
| MT-CYB | 1.46 | 1.63 | 1.59 |
| MT-ATP8 | 1.96 | 1.82 | 1.88 |
| MT-CO1 | 1.90 | 1.76 | 1.74 |
| MT-CO2 | 1.67 | 1.77 | 1.70 |
| MT-ND1 | 1.53 | 1.85 | 1.68 |
| MT-ND2 | 1.62 | 1.80 | 1.64 |
| MT-ND4 | 1.93 | 2.11 | 2.08 |
| MT-ND5 | 1.57 | 1.77 | 1.80 |
| MT-ND6 | 1.66 | 1.78 | 1.74 |
