## Supplementary material for "Hypoxia blunts angiogenic signaling and upregulates the antioxidant system in elephant seal endothelial cells": Table S3

**Table S3. Fold change in expression of respiratory electron chain components in response to long-term hypoxia exposure in human cells.** Fold change is expressed as a comparison to species baseline.

| Gene name | 120 min | 240 min | 360 min |
|---|---|---|---|
| DLAT | 1.15 | 1.23 | 1.18 |
| LDHA | 1.34 | 1.66 | 2.28 |
| MT-ATP6 | 1.67 | 1.84 | 1.62 |
| MT-ATP8 | 1.99 | 1.84 | 1.97 |
| MT-CO1 | 1.87 | 1.83 | 2.08 |
| MT-CO2 | 1.64 | 1.61 | 1.69 |
| MT-CO3 | 1.86 | 1.84 | 1.90 |
| MT-CYB | 1.67 | 1.75 | 1.58 |
| MT-ND1 | 1.70 | 1.80 | 1.53 |
| MT-ND2 | 1.68 | 1.72 | 1.52 |
| MT-ND4 | 2.14 | 2.38 | 2.27 |
| MT-ND5 | 1.79 | 2.02 | 2.00 |
| MT-ND6 | 1.75 | 2.03 | 2.01 |
| PDK1 | 2.24 | 3.27 | 4.60 |
