## Supplementary material for "Hypoxia blunts angiogenic signaling and upregulates the antioxidant system in elephant seal endothelial cells": Table S4

**Table S4. Fold change in expression of TGF-β signaling components in response to 6 h exposure in seal cells.** Fold change is expressed as a comparison to species baseline.

| **Gene name** | **Fold change** |
|---|---|
| MTMR4 | 2.48 |
| NEDD4L | 1.65 |
| SERPINE1 | 1.38 |
| TGFBR1 | 1.36 |
| SMAD2 | 1.28 |
| ITGB1 | 1.13 |
| HDAC1 | 0.81 |
| SMURF1 | 0.76 |
| F11R | 0.71 |
| SMURF2 | 0.66 |
| E2F4 | 0.62 |
| TGFBR2 | 0.58 |
| SMAD3 | 0.55 |
| SKI | 0.54 |
| SKIL | 0.34 |
| TGIF1 | 0.28 |
| JUNB | 0.27 |
| SMAD7 | 0.23 |
