## Supplementary figures and images for "Hypoxia blunts angiogenic signaling and upregulates the antioxidant system in elephant seal endothelial cells"

### Figure S1

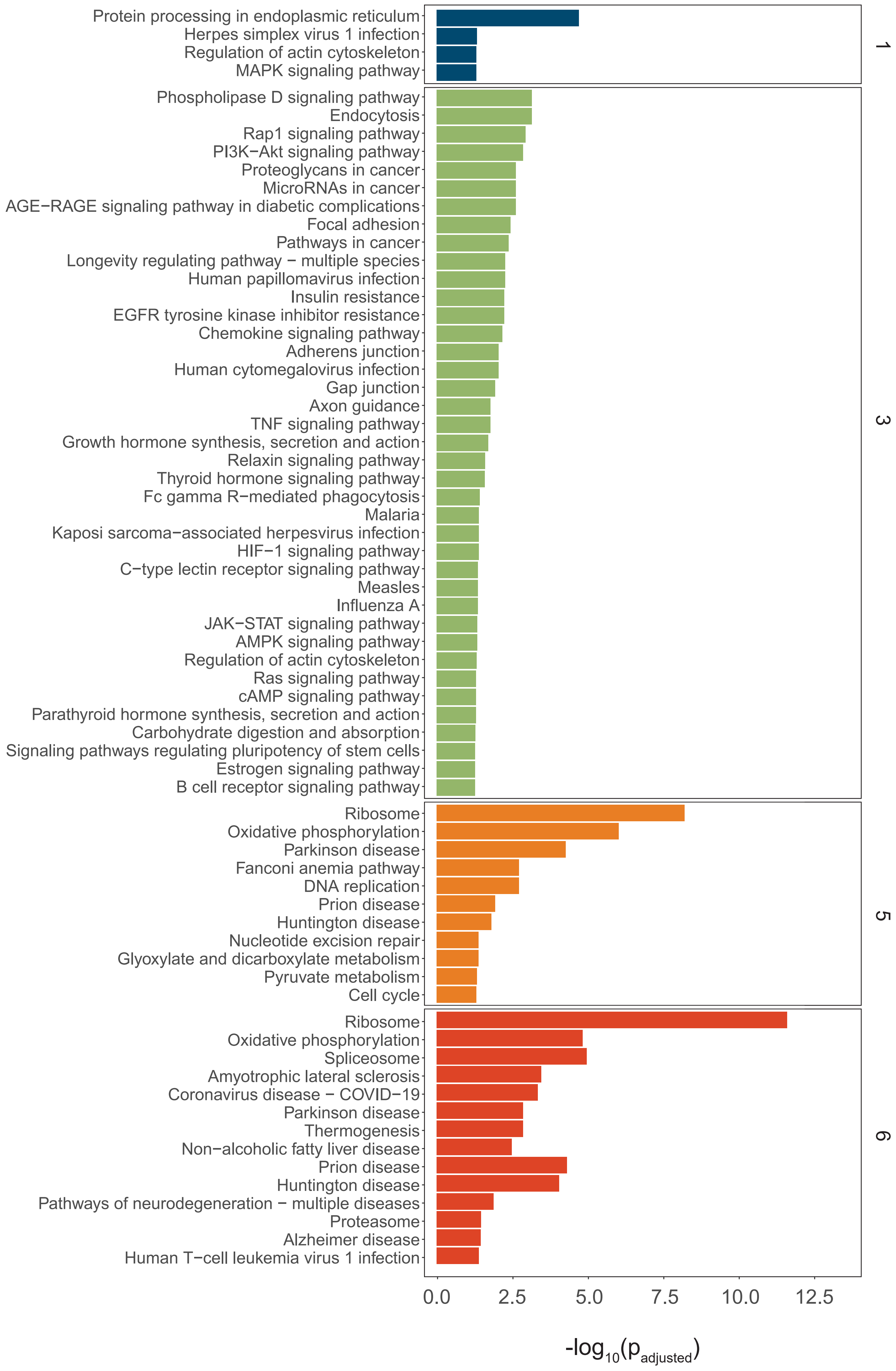

### Figure S2

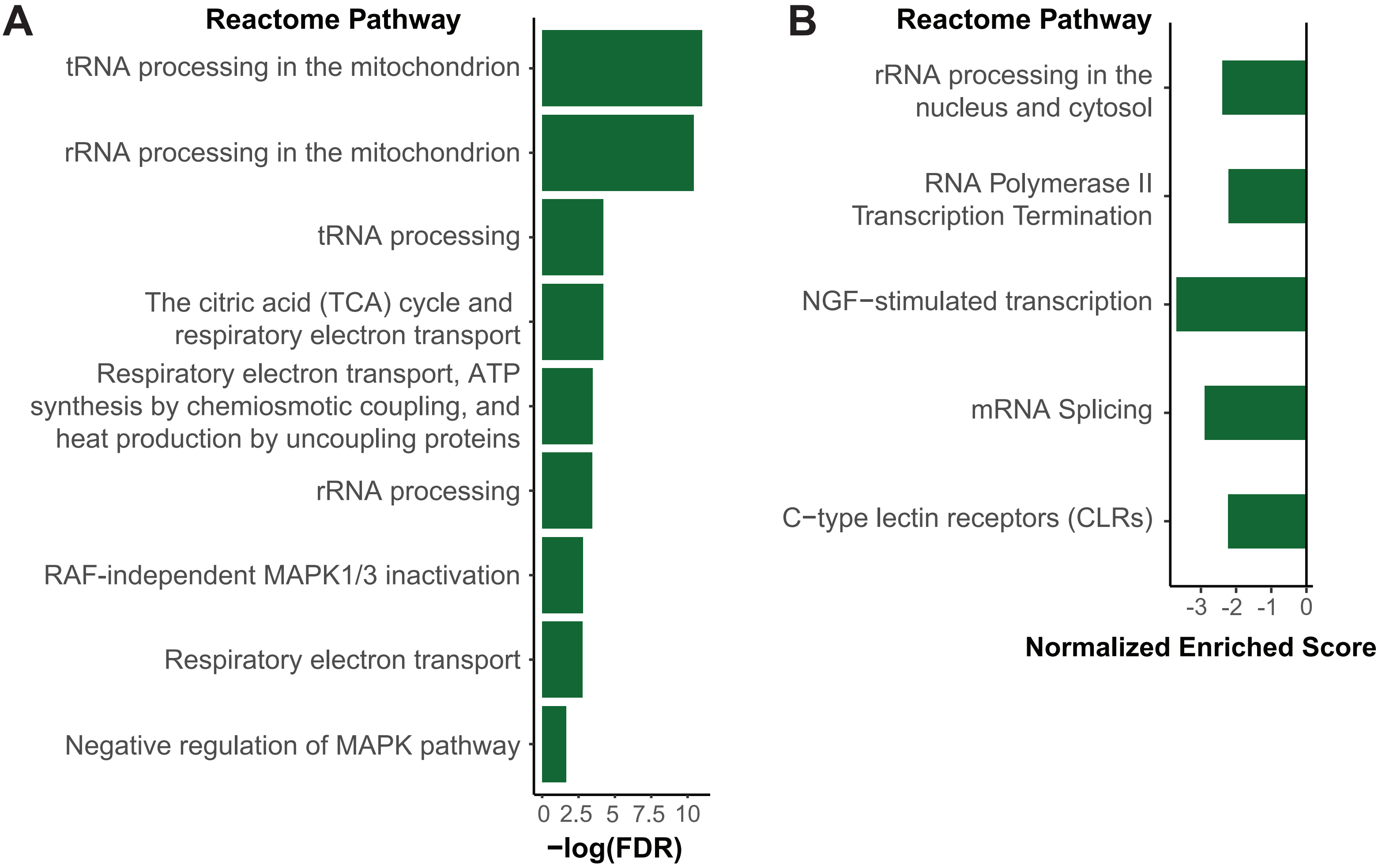
